# Programming dynamic division of labor using horizontal gene transfer

**DOI:** 10.1101/2023.10.03.560696

**Authors:** Grayson S. Hamrick, Rohan Maddamsetti, Hye-In Son, Maggie L. Wilson, Harris M. Davis, Lingchong You

## Abstract

The metabolic engineering of microbes has broad applications, including in biomanufacturing, bioprocessing, and environmental remediation. The introduction of a complex, multi-step pathway often imposes a substantial metabolic burden on the host cell, restraining the accumulation of productive biomass and limiting pathway efficiency. One strategy to alleviate metabolic burden is division of labor (DOL), in which different subpopulations carry out different parts of the pathway and work together to convert a substrate into a final product. However, the maintenance of different engineered subpopulations is challenging due to competition and convoluted inter-strain population dynamics. Through modeling, we show that dynamic division of labor (DDOL) mediated by horizontal gene transfer (HGT) can overcome these limitations and enable the robust maintenance of burdensome, multi-step pathways. We also use plasmid genomics to uncover evidence that DDOL is a strategy utilized by natural microbial communities. Our work suggests that bioengineers can harness HGT to stabilize synthetic metabolic pathways in microbial communities, enabling the development of robust engineered systems for deployment in a variety of contexts.

## Introduction

Many small molecules of commercial, pharmaceutical, or industrial relevance are the products of natural metabolic pathways found in plants, animals, and microbes *(1-5)*. Yields of these pathways of interest are often low, as bulk output is rarely the primary goal for the source organism *(6)*. Many of these pathways can be redirected, optimized, or reengineered for increased output through metabolic engineering *(7-10)*. These systems, with applications in biomanufacturing, bioprocessing, and environmental bioremediation, have the potential to be more cost effective and environmentally friendly than traditional chemical engineering processes *(11-22)*. However, the engineering of complex metabolic pathways often faces several challenges *(23)*.

Optimizing flux through complex pathways in clonal microbial hosts is one such difficulty *(24)*. Natural pathways can be long, consisting of a series of reactions with many intermediates. For example, biosynthesis of the anticancer drug paclitaxel requires at least 19 different enzyme-catalyzed steps in the Pacific yew tree (*Taxus brevifolia*), starting from isoprenyl diphosphate and dimethylallyl diphosphate precursors *(25)*. In natural systems, these pathways are tightly regulated and have been tuned through evolution. Implementing these long pathways in microbial hosts can result in crosstalk between segments of the pathway or between the pathway of interest and other native pathways, reducing their efficiency *(26, 27)*.

The burden imposed on host cells is another major limiting factor in the application of complex metabolic pathways in engineered microbes *(28-30)*. Prolonged burden on the host can drive the emergence of mutants that have lost the engineered function *(31)*. To overcome this limitation, division of labor (DOL) can be an effective strategy to distribute metabolic burden across a consortium of microbes *(32-36)*. In “static” DOL (SDOL), the pathway of interest is partitioned into multiple steps, each being carried out by a different subpopulation. Separating the steps of the pathway reduces the burden on each subpopulation, allowing for each subpopulation to commit more resources to its specific task. If the increase in productivity due to specialization and the additional accumulation of productive biomass outweighs the reduced metabolic efficiency due to the need for transport of intermediates between subpopulations, then an SDOL system will outperform the analogous monoculture *(32, 37)*. Indeed, studies have demonstrated SDOL as a viable approach for executing complex functions *(38-43)*. However, stabilizing inter-strain competition often requires the implementation of population ratio controllers to prevent the extinction of members of the consortium *(44-47)*.

Here, we propose to use horizontal gene transfer (HGT) to achieve *dynamic division of labor* (DDOL) for effective community-based metabolic pathway engineering. HGT has been shown to enhance the functional stability of natural and synthetic microbial consortia by decoupling the persistence of genes and the functions they encode from the abundance of species within the system *(48, 49)*. When fast enough, the transfer of plasmids, or other mobile genetic elements, between microbes overcomes the underlying competitive growth dynamics of subpopulations, stabilizing the function of the population and enabling the persistence of multiple burdensome modules within the system *(50)*. Using kinetic modeling to analyze the population dynamics and productive capacity of monoculture, SDOL, and DDOL systems, we find that DDOL consortia can maintain more burdensome pathways than monoculture systems, while providing superior functional stability compared to SDOL consortia. Through bioinformatic analysis of sequenced plasmids, we also find that DDOL may be a strategy implemented by natural microbial communities to maintain the productivity of some metabolically costly functions, such as symbiotic nitrogen fixation.

## Results

### Quantifying efficiency of a multi-step pathway

We model an *n*-step linear metabolic pathway using a set of coupled ODEs (see Model Formulation). A two-step pathway is illustrated in Figure 1A. The *n*-step pathway converts the substrate (*R*) into a product (*P*) via *n* − 1 intermediate metabolites (*M*_*j*_, *j* = 1, …, *n* − 1). Each step is catalyzed by a single enzyme, *E*_*i*_, with a catalytic constant of *k*_*i*_ and Michaelis-Menten constant of *k*_*i*_. We assume that the substrate, with concentration *R*_*f*_, is fed into the reactor with a dilution rate of *D*.

**Figure 1.**
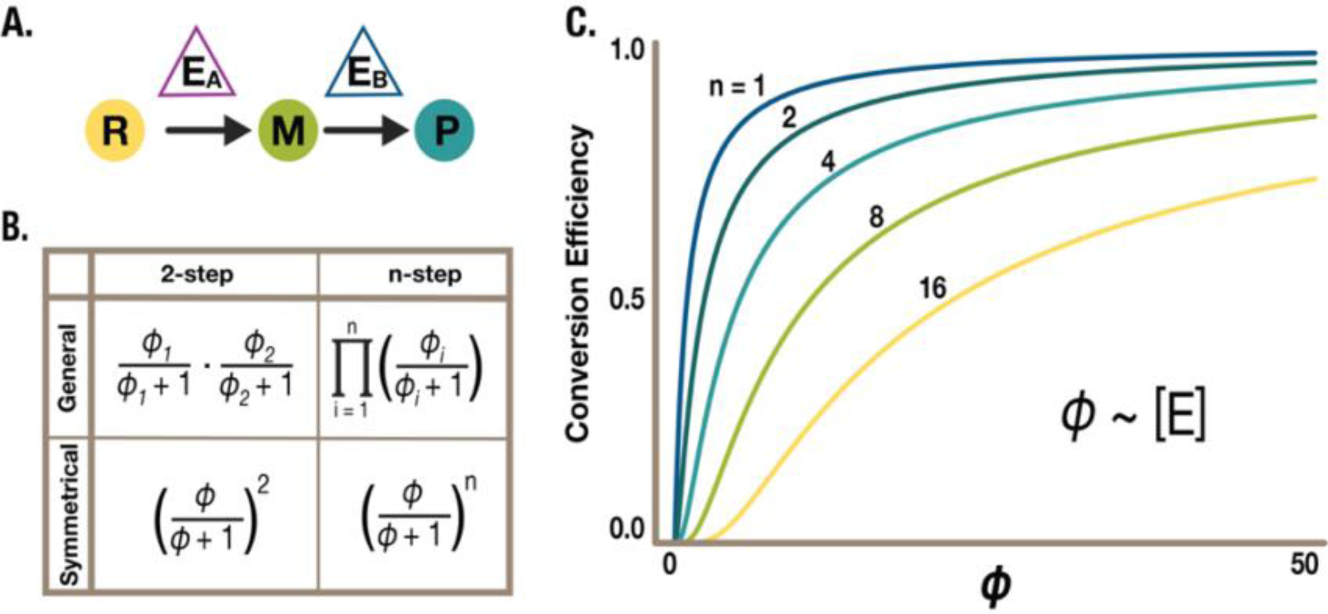
A coarse-grained framework for analyzing pathway activity. A. A two-step metabolic pathway for the conversion of substrate (*R*) to intermediate (*M*) to product (*P*) by two enzymes (*E*_*A*_ and *E*_*B*_) in a single compartment. B. Conversion efficiency of a substrate fed continuously into the reactor (*R*_*f*_) into *P* by the metabolic pathway (*P*/*R*_*f*_). The effective module productivity (*ф*_*i*_) is a function of enzyme concentration (*E*_*i*_), which is in turn proportional to the total biomass of cells producing the enzyme. C. Conversion efficiencies of symmetrical pathways of *n* steps as a function of *ϕ*. The conversion efficiency decreases with the number of steps and, for each pathway, increases with *ϕ*.

At steady state, we have:

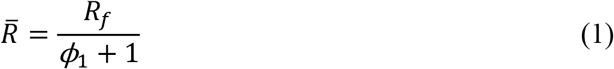

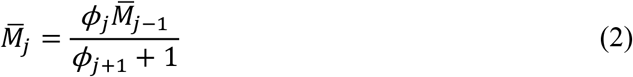

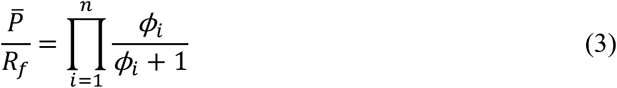

where 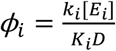 and 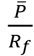 represents the conversion efficiency from the substrate to the product. Equations 12-14 can be further simplified if all steps are equally efficient: *ϕ*_*i*_ = *ϕ*, leading to a symmetric pathway (Figure 1B). As illustrated in Figure 1C, the overall conversion efficiency decreases with *n* but increases with *ϕ*.

Eqs. 1-3 are applicable whether the pathway is carried by a single population or multiple populations. In the latter case, the steady-state solutions are accurate if the enzymatic kinetics operate on a much faster time scale than the population dynamics. Also, *ϕ*_*i*_ will be determined by the total biomass of cells producing *E*_*i*_, which we term the effective biomass. In subsequent analyses, we use effective biomass as our objective function to evaluate the performance of different metabolic pathways.

### DDOL mediated by HGT enables optimal performance of high burden pathways

A two-step pathway can be implemented in a single population via monoculture (Figure 2A), or in multiple populations by SDOL (Figure 2B) or DDOL (Figure 2C). For both monoculture and SDOL implementations, the genes encoding the enzymes cannot move between populations. In DDOL, the genes encoding the enzymes are mobilizable and can transfer between populations. Here we assume the genes are located on plasmids and their transfer is mediated by conjugation, a major mechanism for transfer of diverse functions.

**Figure 2.**
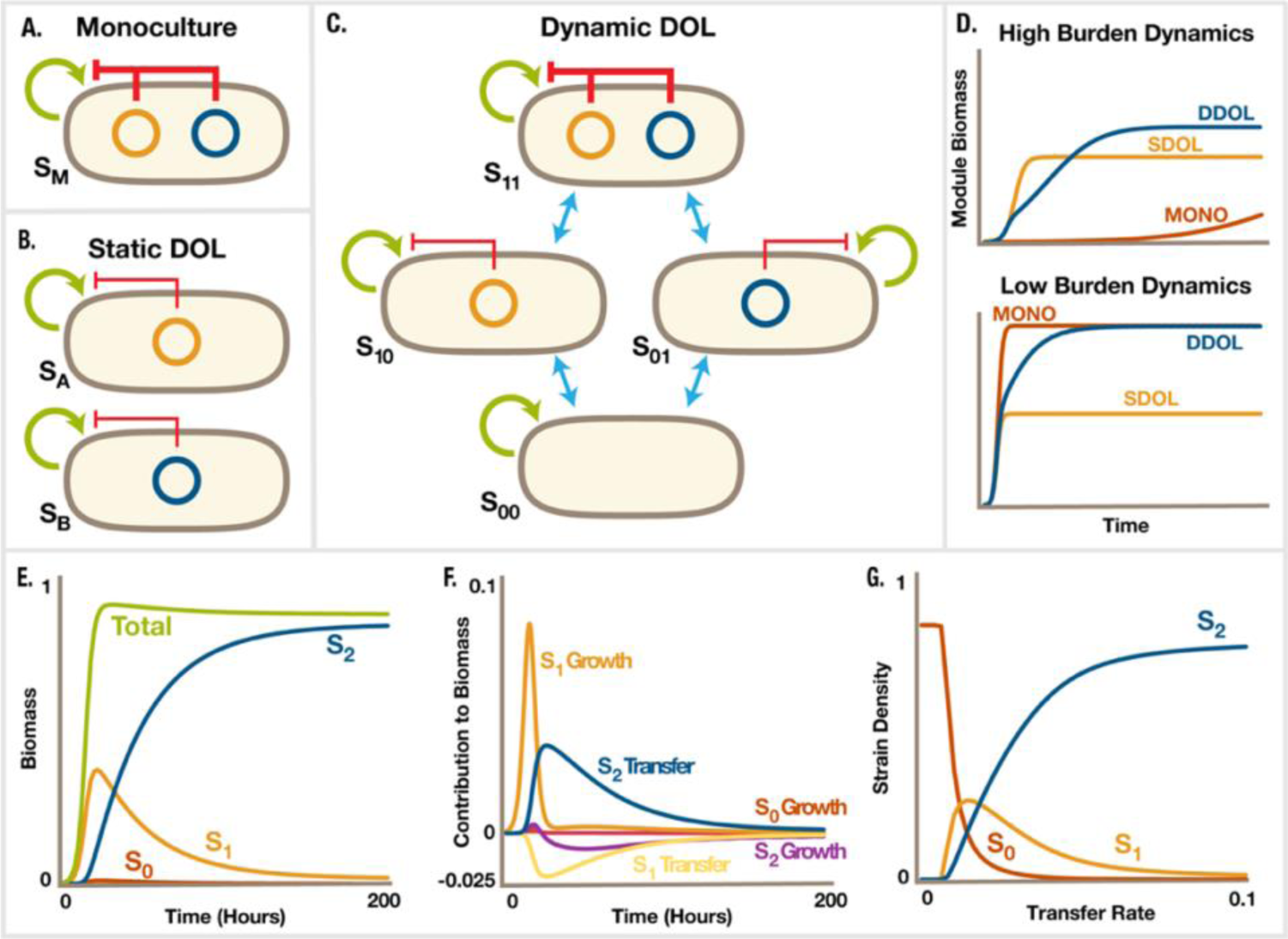
DDOL mediated by HGT enables optimal performance of high-burden pathways. A) A two-step pathway in one population. B) The pathway implemented by SDOL, where two modules are carried by two different populations. Neither module can transfer between populations. C) The pathway implemented by DDOL, where two modules can transfer between populations. D) Representative dynamics of module biomass for each architecture (*m* = 3, *η*_*A*_ = *η*_*B*_ = 0.1, *d*_*A*_ = *d*_*B*_ = 0.001, *D* = 0.05, normalized to maximum growth rate, *μ* _*max*_ = 1). When overall pathway burden is high (*λ*_*A*_ + *λ*_*B*_ = 2.2), DDOL and SDOL systems outperform monoculture systems, while monoculture is more productive when total pathway burden is low (*λ*_*A*_ + *λ*_*B*_ = 0.5). E) Population trajectories for the DDOL system, where S_*i*_ is the curve for a strain carrying *I* modules. For the two-plasmid DDOL system, S_0_ = S_00_, S_1_ = S_10_ = S_01_, and S_2_ = S_11_ (*m* = 3, *η*_*A*_ = *η*_*B*_ = 0.1, *d*_*A*_ = *d*_*B*_ = 0.001, *D* = 0.05, *μ* _*max*_ = 1, *λ*_*A*_ = *λ*_*B*_ = 0.5). F) Contributions of growth and transfer terms to population curves (*m* = 3, *η*_*A*_ = *η*_*B*_ = 0.1, *d*_*A*_ = *d*_*B*_ = 0.001, *D* = 0.05, *μ*_*max*_ = 1, *λ*_*A*_ = *λ*_*B*_ = 0.5). G) Steady state strain densities for a two-module system over varying plasmid transfer rates (*m* = 3, *d*_*A*_ = *d*_*B*_ = 0.05, *D* = 0.05, *μ*_*max*_ = 1, *λ*_*A*_ = *λ*_*B*_ = 0.5).

We further assume that expression of one or both enzymes lead to a growth effect on the host, described by the following equation:

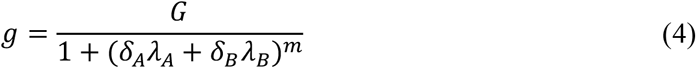

where *δ*_*i*_ = 1 when the host is carrying the genes encoding enzyme *E*_*i*_ and *δ*_*i*_ = 0 otherwise. As the burden felt by the host increases, the effective growth rate, *μ*_*eff*_ = *g μ*_*max*_, decreases. We assume that the burden increases nonlinearly (*m* > 1) with the total amount of each enzyme a cell produces. Under this condition, distributing enzymes between populations can lead to a reduction of the net-burden on the host. Also, we do not consider the potential loss of efficiency in enzymes when they are distributed among multiple populations *(32)*. Considering this factor does not change the qualitative aspects of our conclusions.

When burden is high, DDOL and SDOL outperform the monoculture configuration, while DDOL and monoculture perform similarly when burden is low (Figure 2D). Here we assume that the two enzymes cause equal burden on their host cells. Thus, the two SDOL subpopulations have equal growth rates and can coexist.

For a representative set of parameters (*m* = 3, *η*_*A*_ = *η*_*B*_ = 0.1, *d*_*A*_ = *d*_*B*_ = 0.001, *μ*_*max*_ = 1, *λ*_*A*_ = *λ*_*B*_ = 0.5), the DDOL population dynamics are initially dominated by the growth of the strains carrying only one step (Figure 2E and F). As the total population size approaches the carrying capacity of the system and transfer becomes kinetically favorable, the transfer terms begin to dominate and level the populations out to their steady states. Additionally, the abundance of each plasmid in the DDOL system increases with its transfer rate (Figure 2G). When transfer rates are too low, the plasmids do not persist, and the entire population is made of *S*_00_. As rates of transfer increase, the steady-state populations of *S*_10_ and *S*_01_ increase before being overtaken by *S*_11_ once the transfer rates are sufficient for each plasmid to nearly saturate the system.

### DDOL enables persistence of burdensome, multi-step pathways

HGT allows DDOL systems to maintain more burdensome pathways than SDOL or monoculture (Figure 3). The total burden, *λ*_*T*_,of an *n*-step pathway is given by the following:

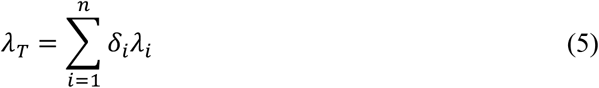

where we set *δ*_*i*_ = 1 for each step of the pathway, *i*. This equation suggests that two factors influence the total burden of a pathway: the burden per step (*λ*_*i*_) and the number of steps (*n*). For a symmetrical, two-step pathway, *λ*_*T*_ = *2λ*_1_. For a representative set of parameters (*m* = 3, *η*_*A*_ = *η*_*B*_ = 0.1, *d*_*A*_ = *d*_*B*_ = 0.001, *D* = 0.02, *μ* _*max*_ = 1), DDOL and monoculture behave similarly when the total burden is low (Figure 3A). As the total burden increases, the monoculture population crashes. Meanwhile, the population density of DDOL systems is maintained, with the productive capacity of the system increasing with transfer rate.

**Figure 3.**
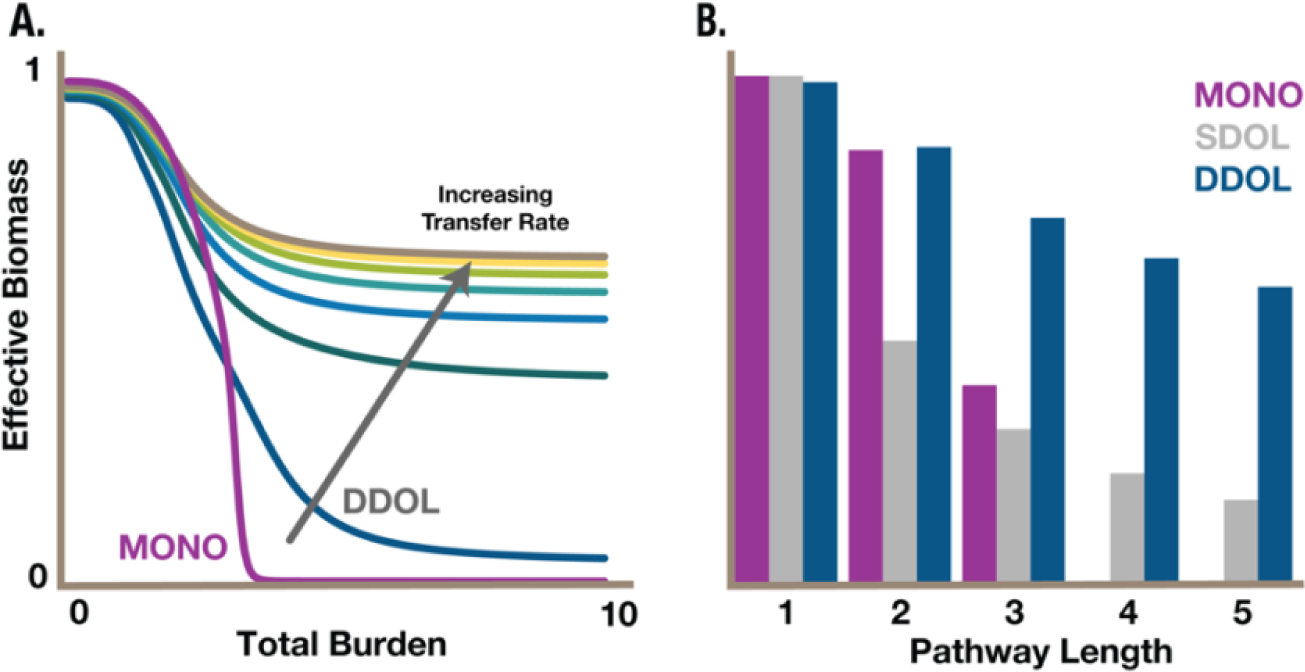
DDOL consortia support high burden and multi-step pathways. A) Effective biomass for a symmetrical, two-step pathway in a monoculture architecture (red curve) and a DDOL architecture (others) with representative parameters (*m* = 3, *η*_*A*_ = *η*_*B*_ ∈ [0.03,0.09], *d*_*A*_ = *d*_*B*_ = 0.001, *D* = 0.02, *μ* _*max*_ = 1). B) Additionally, when burden per step is kept constant and the number of steps increases, DDOL better manages the additional total burden than monoculture and is not as limited by subpopulation carrying capacity as SDOL.

When the burden per step of the pathway is held constant (*λ*_*i*_ = 1), monoculture and DDOL perform similarly for short pathways (Figure 3B). As the pathway length and total burden increase, the monoculture population crashes. DDOL and SDOL are more robust to increasing pathway length and maintain effective biomass for long pathways with high total burden.

### DDOL consortia are more resilient than SDOL consortia to asymmetric burden

Subpopulation dynamics and inter-strain competition can lead to the loss of subpopulations in SDOL communities if they have different growth rates (Figure 4). A comparison of the mean effective biomass for a two-step DDOL pathway and the analogous SDOL pathway shows that the DDOL system is more robust to asymmetric burden (Figure 4A). The difference in mean effective biomass between DDOL and SDOL was calculated by subtracting SDOL quasi-steady-state biomass from DDOL quasi-steady-state biomass for a range of burdens per step. For this representative set of parameters (*m* = 3, *η*_*A*_ = *η*_*B*_ = 0.05, *d*_*A*_ = *d*_*B*_ = 0.001, *D* = 0.03, *μ* _*max*_ = 1), the difference in biomasses ranged from -0.25 to 1. SDOL outperforms DDOL when the burden of the modules is symmetrical, as seen by the yellow streak along the line *y* = *x* in Figure 4A. However, marginal divergences from the line of symmetry result in the loss of one of the SDOL modules and the total loss of pathway productivity (Figure 4B). DDOL is completely insensitive to such divergence.

**Figure 4.**
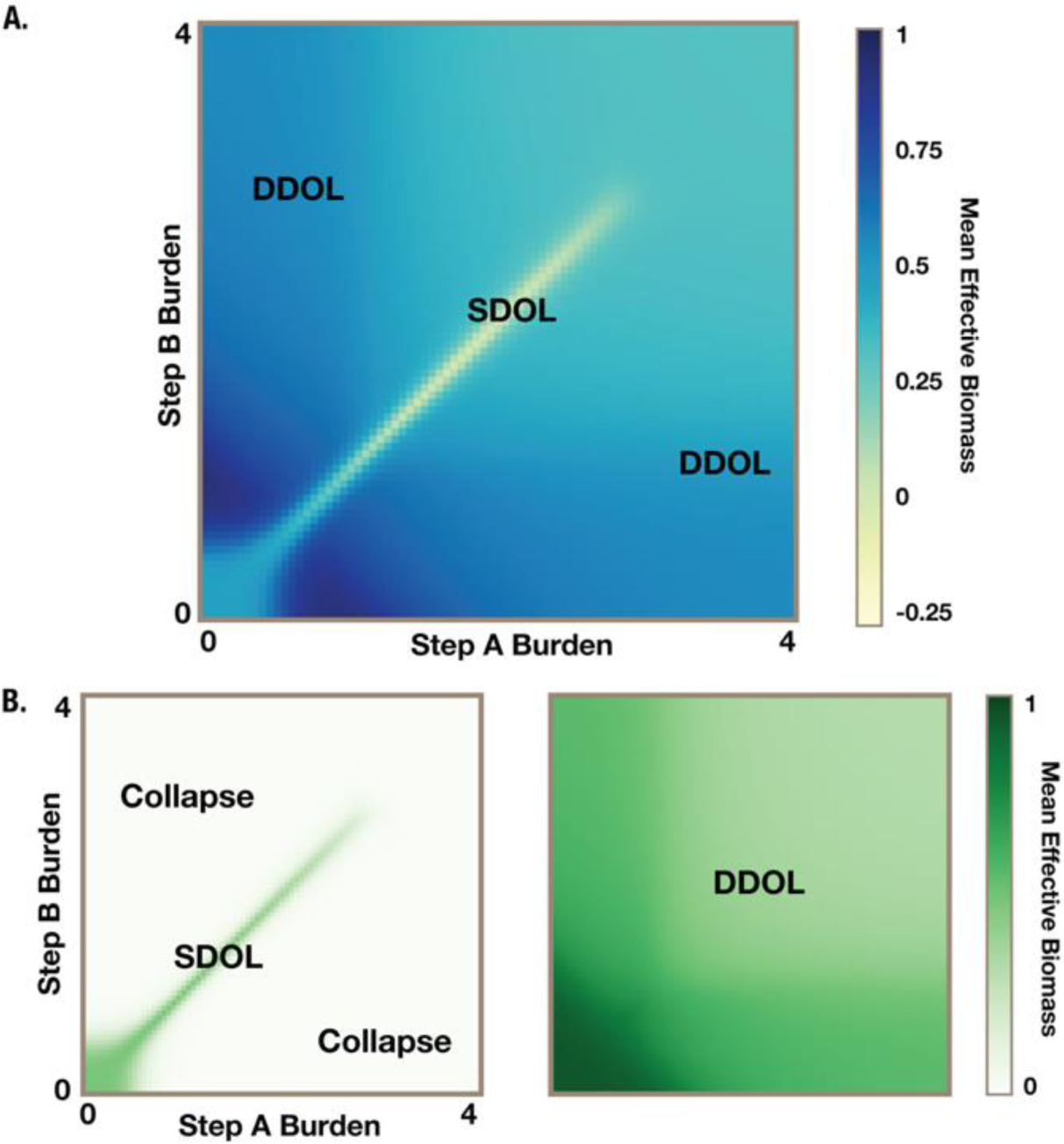
The impact of asymmetric burden on DDOL and SDOL. A) Difference in the geometric mean effective biomass per step between DDOL and SDOL, with SDOL outperforming DDOL along the line *y* = *x* (yellow) and DDOL outperforming SDOL elsewhere (blue). B) Geometric mean effective biomass landscapes for SDOL and DDOL systems ranging from 0 (light green) to 1 (dark green).

### Effective biomass per step can be optimized via transfer and loss rate tuning

Two intrinsic properties of the DDOL system can be tuned such that desired ratios of the steps are obtained: the transfer and loss rates of the plasmids (Figure 5). If the transfer rate of plasmid A is held constant, varying the transfer rate of plasmid B generates a smooth curve of relative steady state abundances for the two plasmids, with the relative abundance of plasmid B increasing with its transfer rate (Figure 5A).

**Figure 5.**
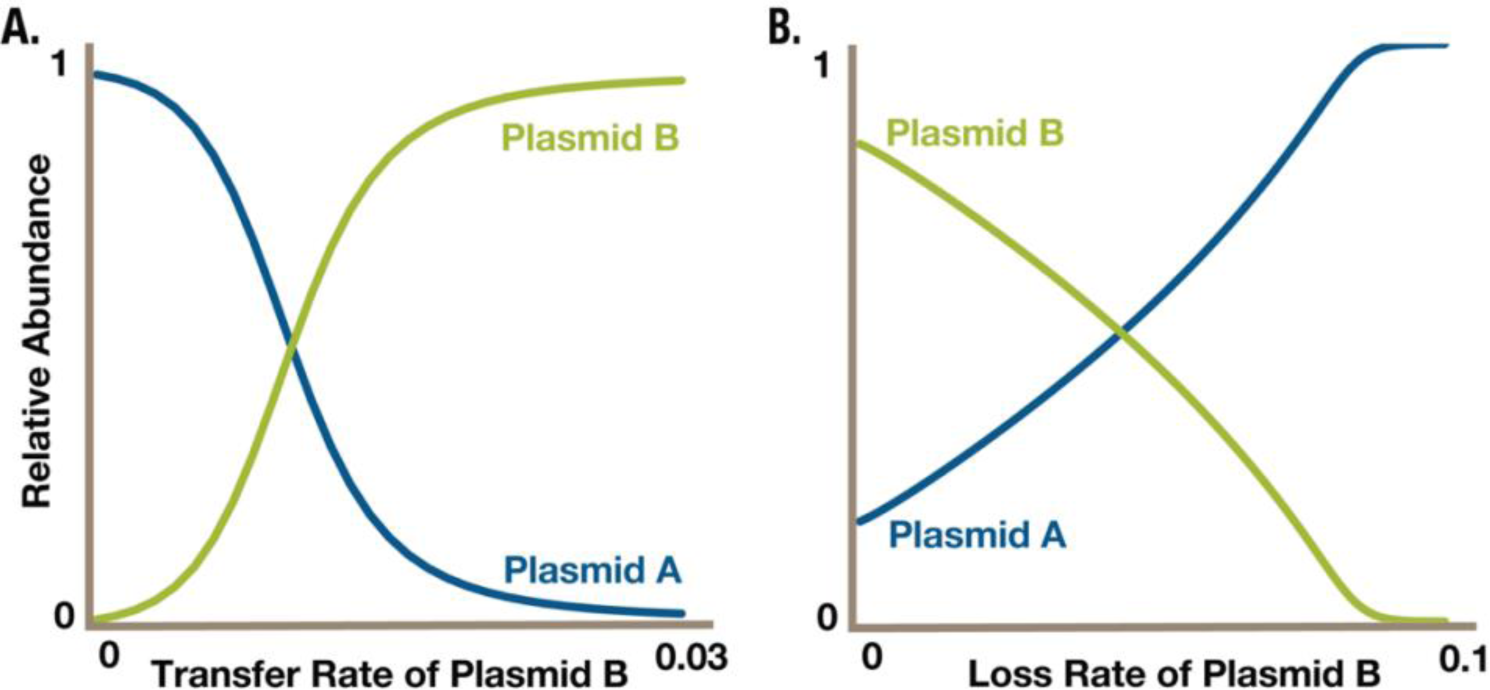
Tuning effective biomass ratios via transfer and plasmid loss rates. A) Increasing the transfer rate of Plasmid B increases its relative abundance in a two-step DDOL system (*m* = 3, *η*_*A*_ = 0.01, *d*_*A*_ = *d*_*B*_ = 0.001, *D* = 0.02, *μ*_*max*_ = 1, *λ*_1_ = *λ*_2_ = 1). B) Choosing a high transfer rate for step B (*η*_*B*_ = 0.1) and low transfer rate for step A (*η*_*A*_ = 0.02) allows for the tuning of module ratios via modifying the loss rate of Plasmid B (*m* = 3, *d*_*A*_ = 0.001, *D* = 0.02, *μ* _*max*_ = 1, *λ*_1_ = *λ*_2_ = 1).

Alternatively, when the transfer rate of plasmid B is high (*η*_*A*_ = 0.1) and the transfer rate of plasmid A is relatively low (*η*_*A*_ = 0.02), the relative abundance of the plasmids can be optimized by tuning the loss rate of plasmid B, such as via induced Cas9 cutting (Figure 5B). When the loss rate of plasmid B is low, plasmid B is more abundant in the system. As the loss rate of plasmid B is increased, the relative abundance of plasmid B declines and the relative abundance of plasmid A increases.

### DDOL is utilized by natural microbial communities

We reasoned that DDOL would be relevant in natural microbial communities in which metabolically burdensome pathways are important for growth and survival. To investigate, we examined the distribution of metabolic genes found on the plasmids in all the complete bacterial genomes in NCBI RefSeq *(51)*. This distribution has a heavy tail (Supporting Figure S1A), and extreme values in this distribution correspond to plasmids found in plant-associated and earth-associated bacteria (Supporting Figure S1B). Many of these bacteria are *Rhizobia* and other diazotrophs like *Azospirillum* that live inside the root nodules of legumes like soybeans (Supporting Figure S1C; Supplementary File S1 with annotations). On average, diazotrophs carry many more metabolic genes on plasmids than non-nitrogen-fixing bacteria (232.6 versus 12.4 metabolic genes on plasmids; Mann-Whitney *U*-test: *p* < 2.2 × 10^-16^; Supporting Figure S2). Gene expression of the *nif/fix* genes (encoding nitrogenases and other genes involved in nitrogen fixation) and *nod* genes (encoding nodulation factors) is metabolically costly, and these genes are often found on symbiotic integrative and conjugative elements (ICEs, also called conjugative transposons) *(52, 53)*. For this reason, we examined the distribution of *nif/fix* and *nod* genes on plasmids. While *nif/fix* genes are found across many bacteria, *nod* genes are almost exclusively found on plasmids containing many *nif/fix* genes (Supporting Figure S3A). We found 444 of these symbiosis plasmids encoding large numbers of both *nif/fix* and *nod* genes. Of these 444 plasmids, 372 are predicted to be conjugative, 24 are predicted to be mobilizable, and 48 are predicted to be non-mobilizable (Supporting Figure S3B). When we compare the mobility of symbiosis plasmids to the mobility of all other plasmids found in their genomes, we find that symbiosis plasmids are far more likely to be conjugative (Figure 6A; Binomial test: *p* < 2.2 × 10^-16^).

**Figure 6.**
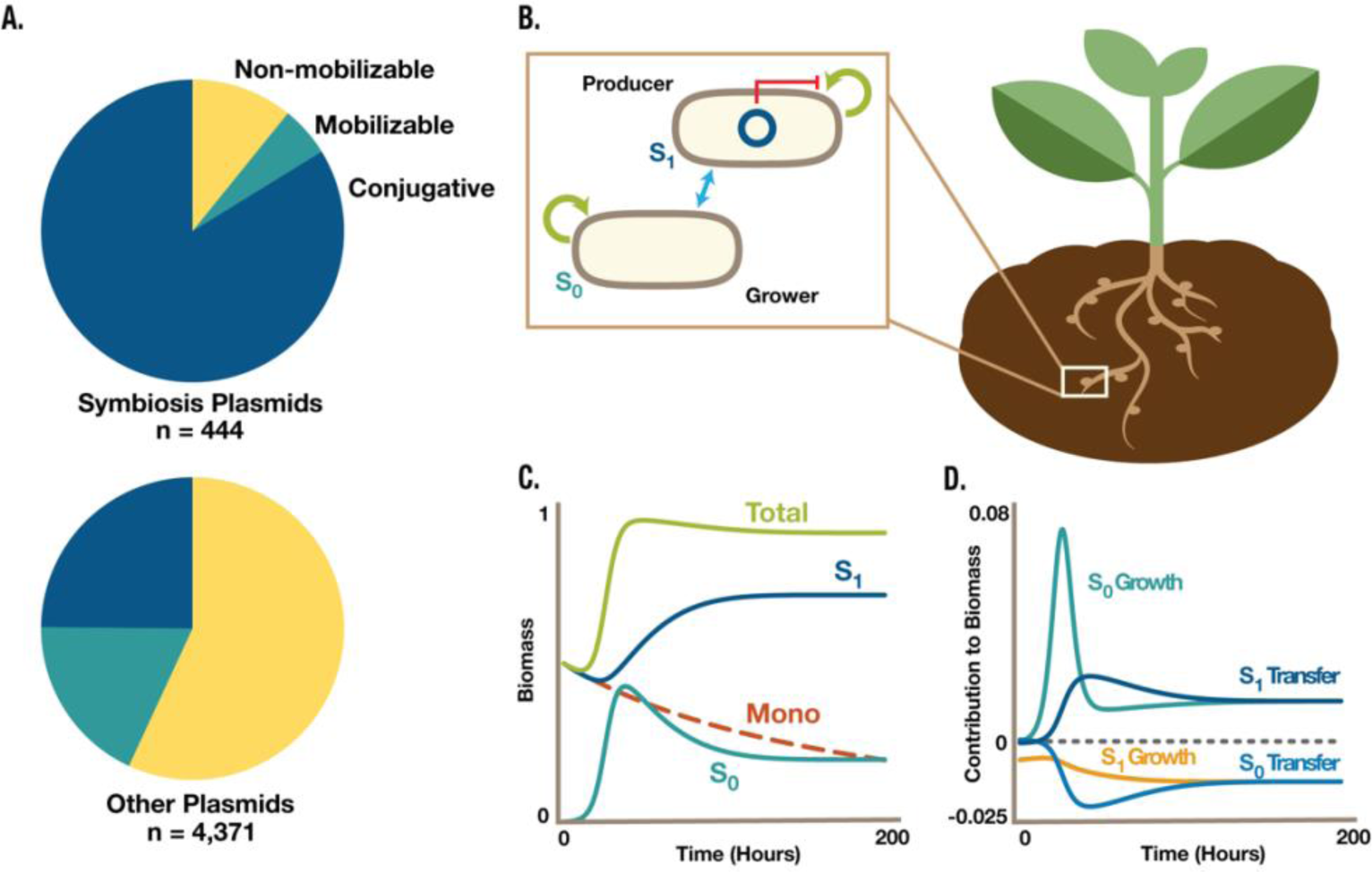
The maintenance of nitrogen-fixing genes via HGT fits the theory of DDOL. A) Distributions of symbiosis plasmids and non-symbiosis plasmids among mobilizability groups. B) Genes for nitrogen fixation are often co-located on a conjugative plasmid in plant-associated microbes. There can be two subpopulations: S_1_, carrying the plasmid, and S_0_, not carrying the plasmid. The transfer and loss of the plasmid mediates flux between the two subpopulations (blue double-sided arrow). C) For a representative set of parameters, a DDOL nitrogen-fixing consortia persists over time, while the analogous monoculture declines in biomass (*m* = 3, *η* = 0.1, *d* = 0.001, *D* = 0.02, *μ* _*max*_ = 1, *λ* = 4, DDOL initial conditions: *S*_0_ = 0, *S*_1_ = 0.5, monoculture initial conditions: *S*_*M*_ = 0.5). D) Growth maintains *S*_0_ biomass, while transfer maintains *S*_1_ by converting *S*_0_ to *S*_1_.

Many metabolic genes in the nitrogen fixation and nodulation pathways are co-localized on single plasmids. In these systems, there are two subpopulations, where one subpopulation carries the plasmid and the other does not (Figure 6B). The transfer and loss of the plasmid allow for the flow of biomass from one strain to the other. After the inoculation of *S*_1_, loss of the plasmid generates a starting population of *S*_0_, which grows rapidly relative to *S*_1_ (Figure 6C). As the total biomass approaches the carrying capacity, HGT becomes more favorable and the *S*_0_ population shrinks as it is converted into *S*_1_. Due to the high burden of the plasmid, *S*_1_ does not grow quickly enough to overcome dilution, but the transfer of the plasmid from *S*_1_ into formerly *S*_0_ cells is sufficient to sustain the *S*_1_ population (Figure 6D). In contrast, the corresponding monoculture population is cannot persist and its biomass crashes (Figure 6C).

## Discussion

We envision DDOL to have several critical advantages over previous approaches for implementing long, burdensome, or complex metabolic pathways in natural and engineered microbial hosts. The burden of implementing heterologous metabolic pathways is often a limiting factor in efficient metabolic engineering for processes like biomanufacturing, biodegradation, and waste remediation. Host cells in a monoculture system are subject to the full burden of the pathway. The persistence of such systems in a continuous flow bioreactor is dependent on the ratio of the dilution rate to the effective growth rate of the burdened strain. Hence, as the total burden of the pathway increases, the growth effect on the monoculture system increases and the monoculture gets washed out of the bioreactor. DDOL systems, however, are not subject to the full brunt of the metabolic burden imposed by the heterologous pathway. Our quantitative theory for DDOL shows that, instead, a small fraction of the population is without plasmid, or with few plasmids, and sustains the total biomass of the population, while conjugation-mediated transfer of the plasmids sustains effective biomass. Due to this balance between growth and productivity, DDOL consortia can support more burdensome pathways than monocultures. This strategy appears to be utilized by natural microbial consortia of great interest, such as *Rhizobia* carrying out the metabolically burdensome processes of plant-associated nodulation and nitrogen fixation.

Recent work in our lab has shown that the transfer dynamics of mobilizable plasmids in a community are more easily controlled than the growth and competition dynamics of the community’s underlying subpopulations *(48)*. Encoding pathway steps on plasmids capable of HGT will therefore allow for the robust programming of the distribution of the pathway within the community, providing for a simpler mode of optimization of the pathway. Initially, as transfer rates increase, DDOL plasmids become more abundant, and the effective biomass of the system increases. However, when transfer rates are too great, the DDOL consortia begins to resemble a monoculture, as plasmids transfer so quickly that they infect every cell. In this case, the effective biomass of a DDOL system follows a similar trend as that of the analogous monoculture, and the DDOL system is unable to sustain high burden pathways. Hence, for the steps of a given pathway, there are optimal transfer rates that can be discovered through experiment-informed modeling and the design-build-test-learn (DBTL) cycle.

Also, metabolic pathways encoded on transferable plasmids are easy to engineer, relative to chromosomal integration, and resilient to deleterious mutations that cause a loss of function within the system *(31)*. The ability to rapidly engineer plasmids accelerates the overall DBTL cycle, and the genetic stability conferred by HGT is a desirable feature for the long-term implementation of microbial consortia to reliably perform robust functions. While the use of plasmids is often avoided for long-term metabolic engineering due to their propensity for loss, transferable plasmids can be maintained within microbial communities in the absence of external selection *(54, 55)*. This allows for the stable persistence of high-copy genes in the system without the need for additional engineering, such as chromosomal tandem gene duplication *(56-58)*.

A recent study has shown that metabolic redundancy within a DOL system can be beneficial by balancing the tradeoffs between metabolic specialization (such as in SDOL) and generality (such as in monoculture) *(59)*. Wang et al. engineered 1,456 unique microbial consortia for the degradation of naphthalene. Each consortium was constructed of strains carrying parts of the naphthalene degradation pathway, with each part of the pathway covered by at least one strain. The authors found that 74 of these consortia outperformed both the monoculture and SDOL versions of the pathway, indicating that some measure of metabolic redundancy can be beneficial for optimizing the productivity of the pathway. Evolving DDOL systems could be one method for identifying the optimal distribution of pathway steps for achieving the ideal degree of metabolic redundancy.

The insight presented here–that the dynamics of burdensome genes in microbial communities can be stabilized via HGT–is not limited to the case of engineering linear metabolic pathways. The stable persistence of many burdensome modules within engineered microbial systems could be used to implement many kinds of complex gene circuits, including those designed for multiplexed biosensing, biocomputing, microbiome engineering, and pattern formation *(60-65)*.

## Methods

### Modeling metabolic pathway flux

We formulated a minimal metabolic pathway model using ordinary differential equations (ODEs) for the concentrations of the small molecule substrate, intermediates, and product of a linear metabolic pathway. We assume that the passive diffusion of small molecules is sufficiently fast that we can approximate the system as one well-mixed compartment.

Consider a simple metabolic pathway entailing the conversion of a substrate (*R*) into an intermediate (*M*) and finally a product (*P*) catalyzed in series by the enzymes *E*_*A*_ and *E*_*B*_. Then the Michaelis-Menten kinetics of the pathway are given by:

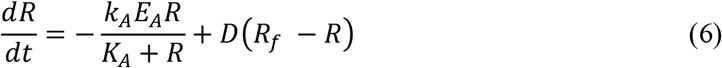

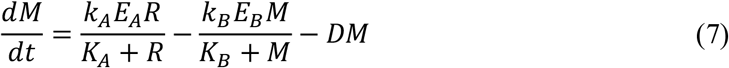

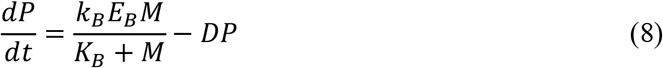

where *R, M*, and *P* are the concentrations of the small molecules in the compartment, *R*_*f*_ is the concentration of the substrate being pumped into the system by the continuous flow bioreactor, *E*_*A*_ and *E*_*B*_ are the concentrations of the enzymes within the compartment, *k*_*A*_ and *k*_*B*_ are the reaction rate constants for the conversion reaction in the pathway, and *k*_*A*_ and *k*_*B*_ are the Michaels-Menten constants for *E*_*A*_ and *E*_*B*_. We assume that *E*_*A*_ and *E*_*B*_ are present at steady state in each cell containing their respective plasmid, so their concentrations are directly proportional to the Step 1 and Step 2 effective biomass, respectively.

### Modeling population dynamics

We modeled the population dynamics of several configurations of microbial factories: monoculture, SDOL, and DDOL. For each population model, we assume that all members of the community follow logistic growth and occupy the same niche. We also assume a continuous dilution term, such as if the microbial cell factory resides within a continuous flow bioreactor with constant influx and efflux of media. Here we describe the dimensionless forms of the population models, normalized to the maximum growth rate of the cell strain (per unit time) and the carrying capacity of the bioreactor (number of cells) where applicable.

We represent the growth dynamics of the monoculture configuration of the microbial cell factory for a two-step metabolic pathway as a single ODE:

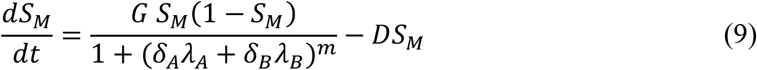

where *S*_*M*_ is the cell density of the monoculture strain, *G* is a catch-all term for general growth rate effects, *λ*_*A*_ and *λ*_*B*_ are the metabolic burden of *E*_*A*_ and *E*_*B*_, respectively, *m* is the Hill coefficient of burden, and *D* is the dilution rate of the bioreactor. For the monoculture, we assume *δ*_*i*_ = 1 for each plasmid.

The analogous SDOL consortium can be described by two ODEs:

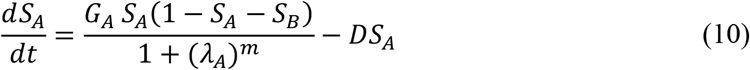

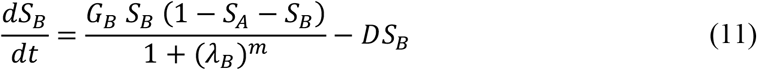

where *S*_*A*_ and *S*_*B*_ are the cell densities of the two strains carrying out complementary steps of the pathway. In the SDOL case, we assume that the modules are not mobilizable and are equally selected for. Additionally, *S*_*A*_ and *S*_*B*_ share the same ecological niche and therefore are subject to the same carrying capacity. By extension, the general ODE for a strain, *S*_*i*_, within a symmetrical *n*-member SDOL consortium is the following:

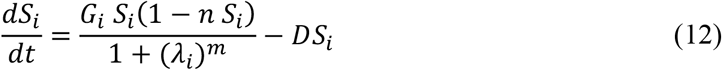

where *δ*_*i*_ = 1 and otherwise *δ*_*j*_ = 0 for *j* ∈ [1, *n*] and *j* ≠ *i*.

Dynamic consortia with transferable plasmids require 2^*n*^ ODEs to model, where *n* is the number of plasmids in the system. We assume that the conjugation of the plasmids does not incur an additional metabolic burden on the host cells *(66)*. For the two-module case, we can model the DDOL community with four ODEs:

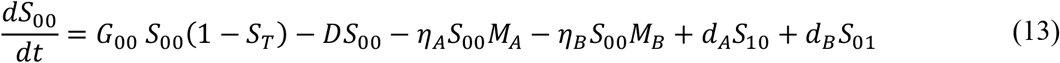

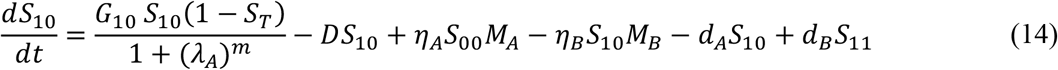

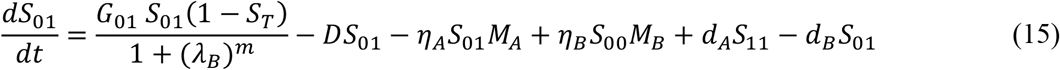

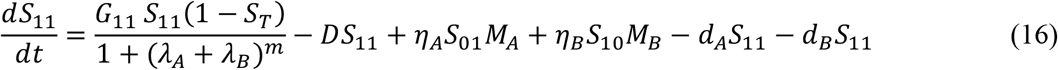

where *S*_00_ is the empty strain (*δ*_*A*_ = *δ*_*B*_ = 0), *S*_10_ carries only plasmid A encoding the first step of the pathway (*δ*_*A*_ = 1, *δ*_*B*_ = 0), *S*_01_ carries only plasmid B (*δ*_*A*_ = 0, *δ*_*B*_ = 1), and *S*_11_ carries both plasmids (*δ*_*A*_ = *δ*_*B*_ = 1). We assume that the four strains occupy the same niche, so we define *S*_*T*_ = *S*_00_ + *S*_10_ + *S*_01_ + *S*_11_. The coefficients *η*_*A*_ and *η*_*B*_ are the conjugation rate constants for the plasmid A and plasmid B, respectively, and we model plasmid transfer using two-body mass action between a strain not carrying a given plasmid and the effective biomass (*M*_*A*_ or *M*_*B*_) of that plasmid. The coefficients *d*_*A*_ and *d*_*B*_ are the loss rates of plasmid A and plasmid B. Note that the DDOL model does not assume selection for either of the plasmids.

### Numerical modeling of population dynamics

All calculations and simulations for the modeling of population dynamics were carried out using the DifferentialEquations.jl package and plotted using the Plots.jl package in Julia 1.9.2. Jupyter Notebooks for the generation of all simulation figures are publicly available at: https://github.com/ghamrick34/DDOL-mathematical-modeling.

### Plasmid genomics data acquisition and analysis

A list of complete prokaryote genomes was downloaded from https://ftp.ncbi.nlm.nih.gov/genomes/GENOME_REPORTS/prokaryotes.txt. The list of prokaryote genomes was then filtered for complete bacterial genomes in NCBI RefSeq containing one or more plasmids, using a Python 3.10 script called *filter-genome-reports*.*py*. A total of 15,693 complete bacterial genomes containing plasmids was downloaded using a Python 3.10 script called *fetch-gbk-annotation*.*py*. Once the genomes were downloaded, tables summarizing the sequence accessions per genome and the genome annotation metadata were generated by Python 3.10 scripts called *make-chromosome-plasmid-table*.*py* and *make-gbk-annotation-table*.*py* and *count-proteins-and-replicon-lengths*.*py*. All plasmids were annotated using MOB-typer 3.1.7 using a Python 3.10 script called *write-plasmid-seqs-for-MOB-typer*.*py* to generate input files for MOB-typer *(67)*. MOB-typer was automated with a Python 3.10 script called *run-MOB-typer*.*py*, and this python script was run on the Duke Compute Cluster (DCC) using a sbatch shell script called *run-MOB-typer*.*sh* as follows: “*sbatch run-MOB-typer*.*sh*”. MOB-typer results for each plasmid were combined into a single text file called *combined_mob_typer_results*.*txt* using a Python 3.10 script called *comb_mob_genome_data*.*py*. Duplicate MOB-typer results were identified by running the following one-line shell script: *sort combined_mob_typer_results*.*txt*| *sort* | *uniq -d > duplicated*.*txt*. Then, the number of duplicate entries was counted with this one-line shell script: *cat duplicated*.*txt* | *wc -l*. Then, a final text file without duplicate entries was generated with the following one-line-shell script: *sort -u combined_mob_typer_results*.*txt -o unique_mob_results*.*txt*. Finally, plasmid mobility predictions from MOB-typer were parsed with a Python 3.10 script called *parse-MOBtyper-results*.*py*.

We annotated metabolic genes on plasmids as follows. First, we made a database of all proteins found on plasmids with a Python 3.10 script called *make-plasmid-protein-FASTA-db*.*py*. We then ran a Python 3.10 script called *make-GhostKOALA-input-files*.*py* to generate input files for the GhostKOALA webserver associated with the Kyoto Encyclopedia of Genes and Genomes (KEGG) database *(68-72)*. The GhostKOALA algorithm annotates the function of proteins based on homology to proteins with known functions in the KEGG database *(68)*. Results from the GhostKOALA webserver were downloaded and concatenated with a Python 3.10 script called *concatenate-and-filter-GhostKOALA-results*.*sh*. Additional quality control to remove annotation errors was applied, using a Python 3.10 script called *remove-chromosomes-from-GhostKOALA-results*.*py*. Then, unique KEGG Orthology IDs (each corresponding to a class of protein homologs with a common function) in the set of plasmid proteins were extracted with a Python 3.10 script called *get_unique_KEGG_IDs*.*py*. These KEGG IDs were uploaded to the KEGG Mapper Reconstruct webserver at: https://www.kegg.jp/kegg/mapper/reconstruct.html. Unique plasmid protein KEGG IDs mapping to metabolic pathways in KEGG were downloaded. Finally, a Python 3.10 script called *get-plasmid-metabolic-KOs*.*py* was used to annotate all plasmid proteins mapping to metabolic pathways in the KEGG database.

We used the “host” and “isolation_source” fields in the RefSeq annotation for each genome to place each into the following categories: Marine, Freshwater, Human-impacted (environments), Livestock (domesticated animals), Agriculture (domesticated plants), Food, Humans, Plants, Animals (non-domesticated animals, also including invertebrates, fungi and single-cell eukaryotes), Soil, Sediment (including mud), Terrestrial (non-soil, non-sediment, including environments with extreme salinity, aridity, acidity, or alkalinity), NA (no annotation). These annotations were based on the annotation categories in the ProGenomes2 Database (Aquatic, Disease associated, Food associated, Freshwater, Host associated, Host plant associated, Sediment mud, Soil). The main difference between our annotation categories and those used in the ProGenomes2 database is that our annotations split host-associated categories based on domestication, and bin all human-host associated microbes together, regardless of disease association. For reproducibility, our annotations are generated using a Python 3.10 script called *annotate-ecological-category*.*py*. To simplify the data presentation, we merged categories as follows. Marine and Freshwater categories were grouped as “Water”. Sediment, Soil, and Terrestrial categories were grouped as “Earth”. Plant and Agriculture categories were grouped as “Plants”.

A shell script called *rhizobial-symbiosis-plasmid-protein-grep*.*sh* was used to find all plasmid proteins encoding Nif/Fix nitrogen-fixation genes and Nod root nodulation genes, as well as the genomes containing these plasmids. All figures and analyses of metabolic genes on plasmids, with a focus on Nif/Fix and Nod rhizobia-legume symbiosis pathways, were generated with an R 4.2 script called *make-DDOL-figures*.*R*. All plasmid and genome data used in this bioinformatic analysis is publicly available from the NCBI RefSeq database. All code sufficient to reproduce the bioinformatic analysis is publicly available at: https://github.com/rohanmaddamsetti/DDOL-plasmid-bioinformatics/.

## Acknowledgments

The authors would like to thank Ryan Tsoi, Teng Wang, Emma Chory, Amy Schmid, and Pranam Chatterjee for insightful comments and suggestions, and Tom Milledge and Duke Research Computing for assistance with the Duke Compute Cluster. This work was supported by the National Science Foundation Grant DGE 2139754 (to G.S.H.) and the National Institutes of Health Grants 1T32GM144291 (to G.S.H.) and R01EB031869 (to L.Y.).

## Author Information

### Authors

Grayson S. Hamrick – Department of Biomedical Engineering, Center for Quantitative Biodesign, Center for Biomolecular and Tissue Engineering, Duke University School of Medicine, Durham, North Carolina, 27708, USA

Rohan Maddamsetti – Department of Biomedical Engineering, Center for Quantitative Biodesign, Duke University School of Medicine, Durham, North Carolina, 27708, USA

Hye-In Son – Department of Biomedical Engineering, Center for Quantitative Biodesign, Duke University School of Medicine, Durham, North Carolina, 27708, USA

Maggie L. Wilson – Department of Biomedical Engineering, Center for Quantitative Biodesign, Duke University School of Medicine, Durham, North Carolina, 27708, USA

Harris M. Davis – Department of Biomedical Engineering, Center for Quantitative Biodesign, Duke University School of Medicine, Durham, North Carolina, 27708, USA

### Author Contributions

G.S.H. and L.Y. devised the project. G.S.H. and L.Y. formulated theoretical framework. G.S.H and H.M.D. performed numerical simulations. R.M., M.L.W., and H.S. performed bioinformatic analysis. G.S.H., R.M., and L.Y. wrote the manuscript with input from all authors.

### Competing Interests

The authors declare no competing interests.

### Data Availability

All data is available in the main text and the supporting information, or on request from the corresponding author. Modeling code is available directly at https://github.com/ghamrick34/DDOL-mathematical-modeling. Bioinformatics code is available directly at https://github.com/rohanmaddamsetti/DDOL-plasmid-bioinformatics.

## Notes

https://github.com/ghamrick34/DDOL-mathematical-modeling

### Competing Interest Statement

The authors have declared no competing interest.

