## Supporting Material for "Programming dynamic division of labor using horizontal gene transfer"

**SUPPORTING SECTION 1. SUPPORTING FIGURES AND TABLES 2**

#### Supporting Figures

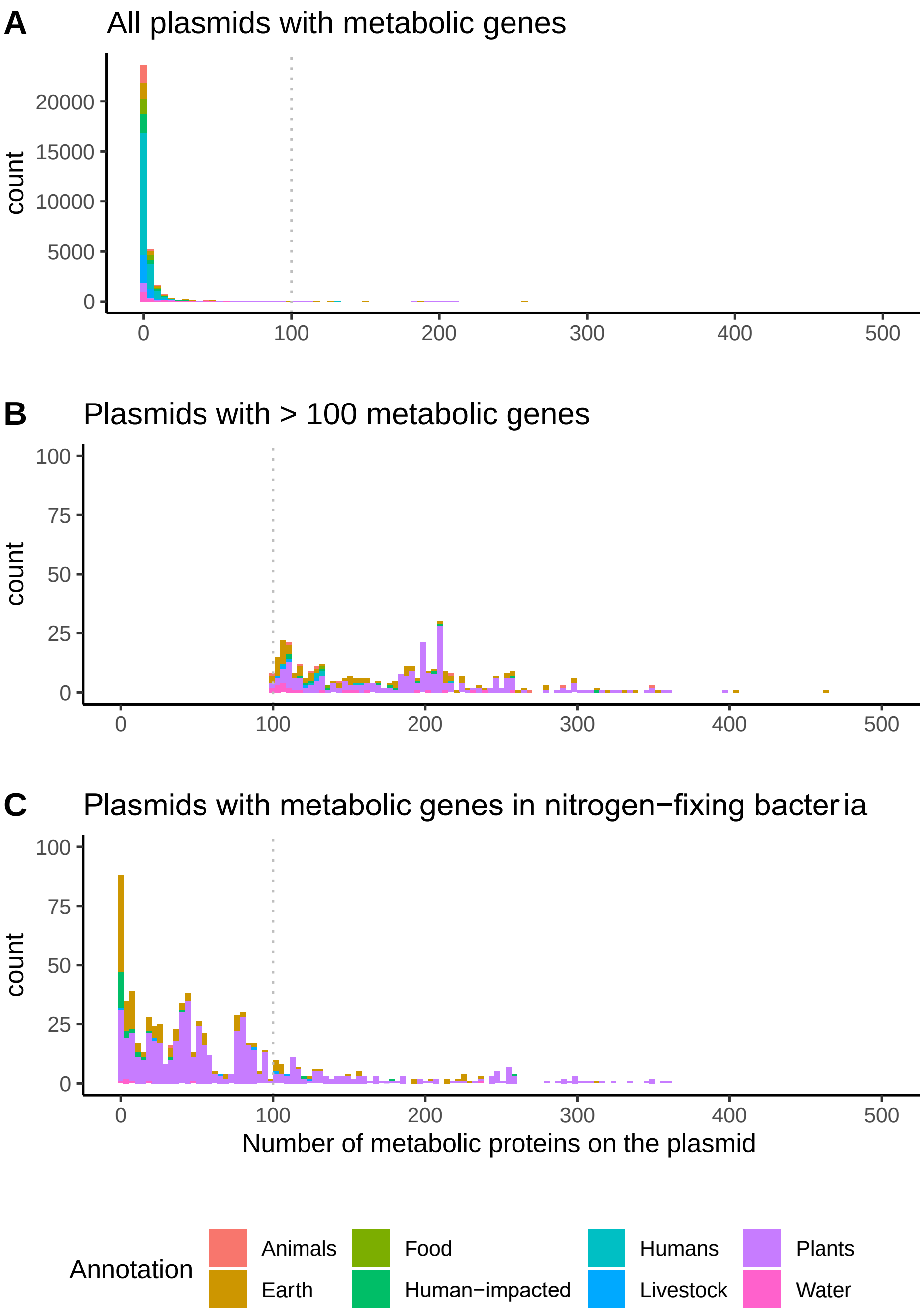


**Supporting Figure S1. Nitrogen-fixing bacteria contain plasmids with hundreds of metabolic genes.**

A) The distribution of plasmids containing metabolic genes is heavy tailed.
B) Extreme values in this distribution correspond to plasmids found in plant-associated and earth-associated bacteria.
C) Nitrogen-fixing bacteria account for a substantial subset of plasmids in the heavy tail.

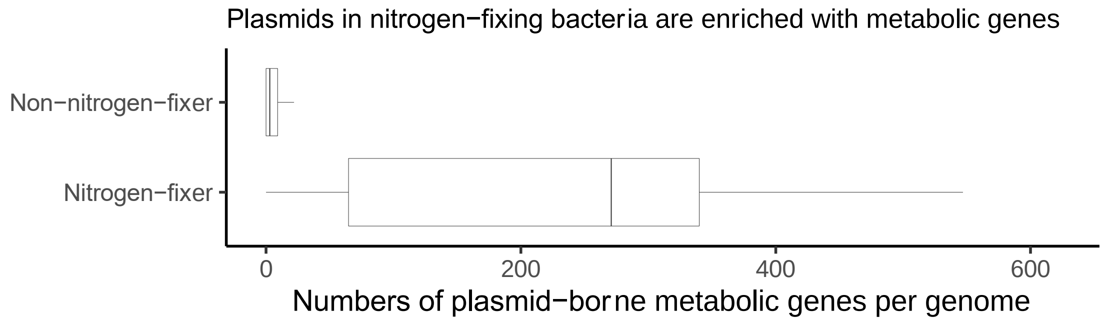


**Supporting Figure S2. On average, nitrogen-fixing bacteria carry more metabolic genes on plasmids than non-nitrogen-fixing bacteria.**

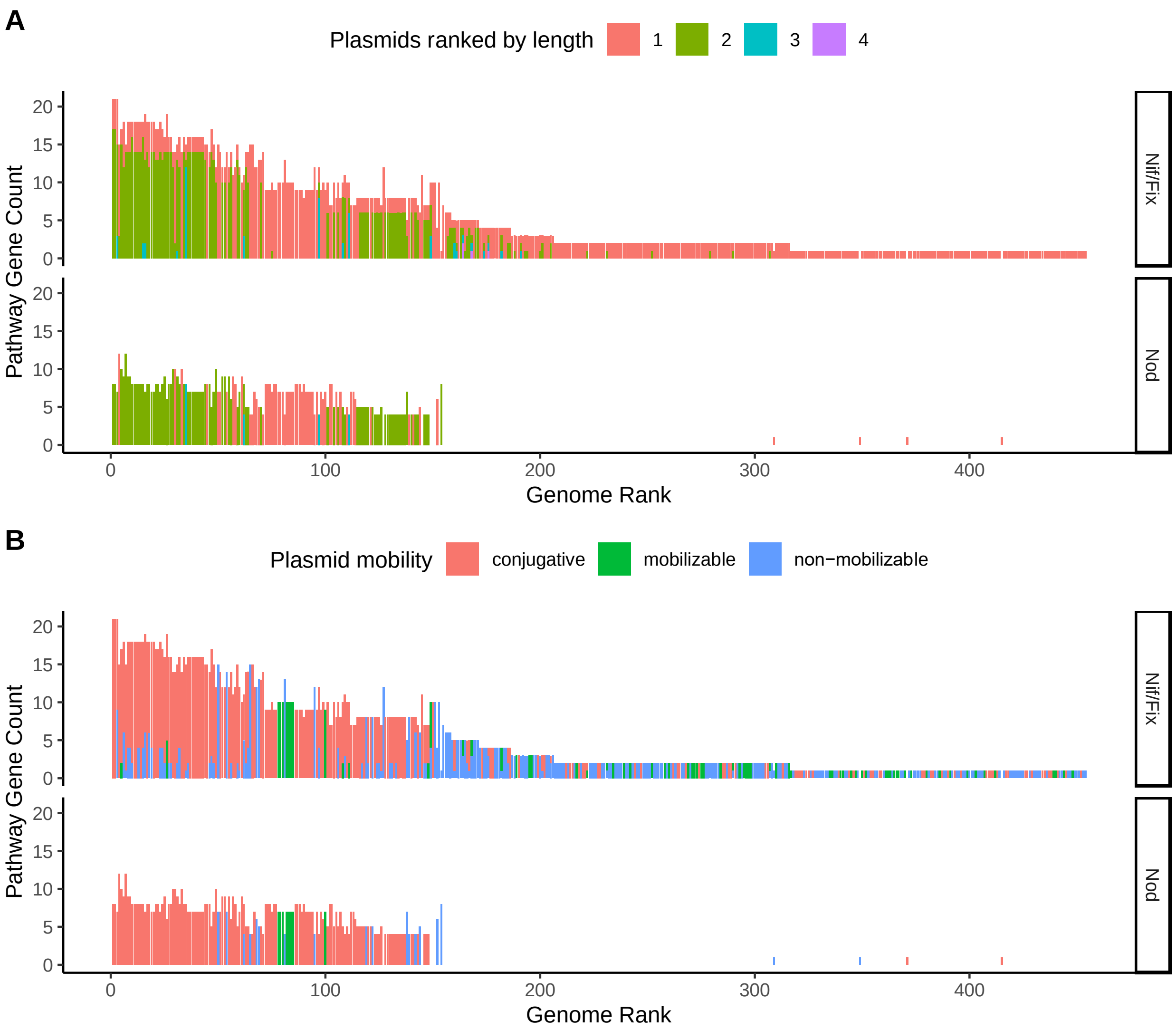


**Supporting Figure S3. Nod genes co-localize with Nif/Fix nitrogen-fixation genes on conjugative symbiosis plasmids.** Genomes are ranked by the number of Nif/Fix and Nod genes on their plasmids. Each bar represents a single genome, and the length of each bar corresponds to the number of genes in either the Nif/Fix or Nod pathway found on plasmids in that genome. Plasmids in each genome were ranked by length.

A) Sections of each bar are colored to indicate the identity of the plasmid containing Nif/Fix or Nod genes in each genome. The length of the red bar corresponds to the number of Nif/Fix or Nod genes found in the largest plasmid containing Nif/Fix or Nod genes in the genome. Green corresponds to the second-largest plasmid containing Nif/Fix or Nod genes, and so on for the third largest (blue) or fourth-largest (purple).

B) Sections of each bar are colored to indicate the mobility of each plasmid containing Nif/Fix or Nod genes in each genome. Red indicates conjugative plasmids, green indicates mobilizable plasmids, and blue indicates non-mobilizable plasmids.

### Supporting Tables

**Supporting Table S3. Parameter values used for numerical simulations.** Modeling parameters were non-dimensionalized (ND) with respect to carrying capacity and maximum growth rate of the unburdened strain.

| Parameter | ND Parameter | Biological Range | ND Range | Base Modeling Value | Reference |
|---|---|---|---|---|---|
| Maximum Growth Rate | $\mu_{\max}$ | 0.5 $hr^{-1}$ | – | 1 | *(32)* |
| Carrying Capacity | $\rho$ | 1 $\mu$L | – | 1 | *(32)* |
| Burden | $\lambda_i$ | – | [0,32] | 1 | *(32)* |
| Growth Effect | $G$ | – | [0,10] | 1 | *(32)* |
| Dilution Rate | $\mathcal{D}$ | 0.05 $hr^{-1}$ | [0,1] | 0.1 | *(55)* |
| Hill coefficient of burden | $m$ | – | [1,10] | 3 | *(32)* |
| Basal Plasmid Loss Rate | $d_i$ | $5 \times 10^{-4}$ $hr^{-1}$ | [0,0.1] | 0.001 | *(55)* |
| Conjugation Efficiency | $\eta_i$ | $[0, 10^{-10}]$ $cell^{-1}$ $hr^{-1}$ | [0,0.2] | 0.05 | *(55, 66)* |
